# Control of helical navigation by three-dimensional flagellar beating

**DOI:** 10.1101/2020.09.27.315606

**Authors:** Dario Cortese, Kirsty Y. Wan

## Abstract

Helical swimming is a ubiquitous strategy for motile cells to generate self-gradients for environmental sensing. The model biflagellate *Chlamydomonas reinhardtii* rotates at a constant 1 – 2 Hz as it swims, but the mechanism is unclear. Here, we show unequivocally that the rolling motion derives from a persistent, non-planar flagellar beat pattern. This is revealed by high-speed imaging and micromanipulation of live cells. We construct a fully-3D model to relate flagellar beating directly to the free-swimming trajectories. For realistic geometries, the model reproduces both the sense and magnitude of the axial rotation of live cells. We show that helical swimming requires further symmetry-breaking between the two flagella. These functional differences underlie all tactic responses, particularly phototaxis. We propose a control strategy by which cells steer towards or away from light by modulating the sign of biflagellar dominance.

---

Organisms perceive their world as three-dimensional. Biological swimmers adopt chiral or helical trajectories to navigate through bulk fluid. Helical navigation is a ubiquitous locomotion strategy found across diverse taxa and multiple length scales, from the planktonic larvae of marine invertebrates [1] to small protists [2] and spermatozoa [3]. Bacteria bias the rotation of chiral flagella in response to chemical gradients [4]. In contrast, many unicellular eukaryotes *steer* towards or away from external stimuli (light, chemicals, gravity) via helical klinotaxis, and a corkscrewing motion around their body axis. Such organisms routinely integrate sensory information obtained by subcellular sensors (receptors, eyespots) which periodically scan the environment, with motor actuators (cilia, flagella), to adjust their swimming trajectories according to the stimulus. These self-actions can enhance signal perception at the microscale [5], with important consequences for evolution and eukaryogenesis [6]. Helical movement could help compensate for asymmetries in body shape [7] and/or filament actuation [8], but it is unclear whether such asymmetries evolved as an adaptation to helical swimming or to facilitate it in the first place. Self-propelled cholesteric liquid crystal droplets also exhibit curling and helical motions, depending on an interplay between surface flows and nematic order [9]. The complex motility strategies of real cells can facilitate the development of next-generation artificial swimmers and controllable devices [10].

For algal flagellates, the control of flagellar beating is key to effective spatial navigation and reorientation to photostimuli [11–13]. Spermatozoa steer via changes to their 3D beat pattern [14], while ciliates tilt the orientation of ciliary fields to turn [15]. *Chlamydomonas* swims with two near-identical flagella using a characteristic breaststroke, rotating slowly (1-2Hz) about its axis as it swims along left-handed helices [16, 17]. Each (~ 10 *μm*) cell has a unique photosensor (eyespot), fixed in position on the rigid cell body [18]. The eyespot scans the environment and modulates phototactic turning by periodic shading [19, 20]. Yet to date, there has been no compelling explanation for the cell’s characteristic helical motion. Here, we combine theory and experiment to reveal how *Chlamydomonas*, despite a very symmetric body plan, manipulates non-planar biflagellar beating to achieve helical swimming and stimulus-dependent steering responses in three dimensions. This is in contrast to other microswimmers which rely on obvious body asymmetries to steer [21], such as two very different types of flagella in the case of dinoflagellates [22].

## Model formulation

We construct a fully-3D model of the swimmer, as three beads interacting hydro-dynamically in an incompressible newtonian fluid at zero-Reynolds number. The cell body is a sphere of radius *a*_0_, located at ***r***_0_. Two smaller beads (radius *a* ≪ *a*_0_) are localised to the approximate centers of drag of the flagella, at ***r***_i_, *i* = 1, 2. Flagella beads are driven by variable tangential forces 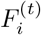, *i* = 1, 2 and constrained to move along circular orbits of radius *R* by normal components 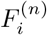, to mimic breaststroke swimming. The 3 beads are constrained on a rigid triangular scaffold assumed to be at rest with the body frame of reference (Fig. 1), with dimensions *ℓ* and *h*, and *a* ≪ *a*_0_ ≪ *h, ℓ*. This is motivated by experiments which show that freely-swimming *Chlamydomonas* cells induce flow-fields that are well-described by just three Stokeslets [23]. In-plane versions of these models recapitulated the stochastic (run-and-tumble) character of biflagellar coordination [24–27]. These minimal representations of ciliary beating as beads moving along a prescribed orbits are powerful tools for studying hydrodynamic synchronization [28–30].

**FIG. 1.**
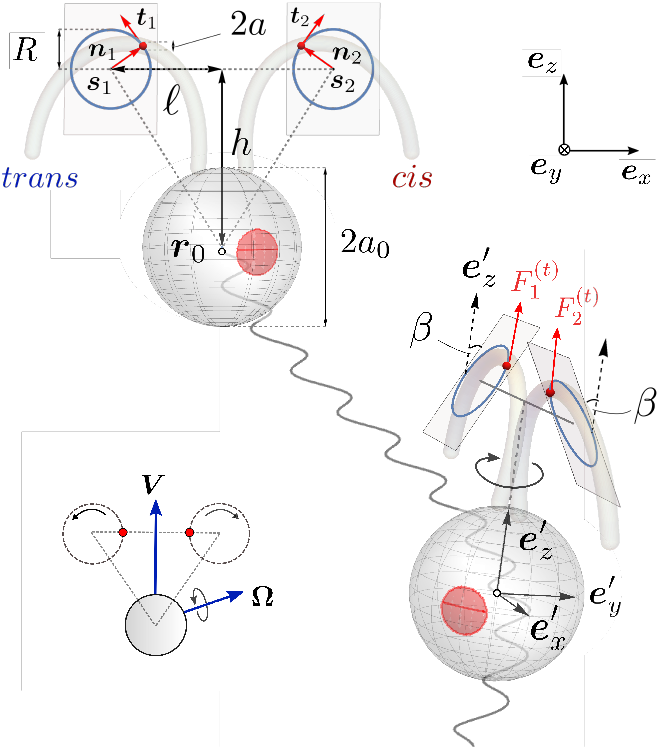
A fully-3D model of a freely-swimming *Chlamydomonas* cell, in front, and side views. Flagellar beating is modelled by small beads constrained to rotate along circular orbits embedded within a pair of tilted beat planes, for a realistic scaffold geometry (inset). 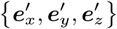 and 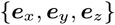 are the body and laboratory reference frames.

Unlike in previous studies, we allow the flagellar beads to *rotate out of the plane*. This introduces a non-zero tilt angle *β* between the flagellar orbital planes *π*_(±*β*)_ and the frontal 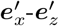 plane, which we hypothesize suffices to produce both axial rotation and helical swimming (Fig. 1b). The flagella orbits are centered at 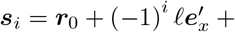 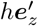, *i* = 1, 2. Flagellar beads, located at ***r***_*i*_ = ***s***_*i*_ + *R**n***_*i*_ and 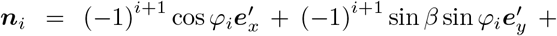 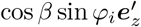, rotate with phases *φ*_*i*_. For breaststrokes, we require 0 ≤ *β* < *π*/2, 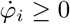. The body axes 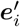 transform to the lab frame ***e***_*i*_ via Euler angles ***θ*** = (*θ*_1_, *θ*_2_, *θ*_3_) (yaw, pitch and roll). The swimmer kinematics are fully described by ***X*** = (*x*_0_, *y*_0_, *z*_0_, *θ*_1_, *θ*_2_, *θ*_3_, *φ*_1_, *φ*_2_) and the parameters *h, ℓ, β, R, a*_0_, *a*. We impose force- and torque-free conditions: ***F***_0_ + ***F***_1_ + ***F***_2_ = **0** and ***T***_0_ + ***T***_1_ + ***T***_2_ = **0**, where 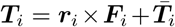, and 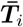 is the intrinsic torque due to the *i*th sphere’s rotation around an internal axis. Since *a* ≪ *a*_0_, we assume that 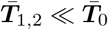. The swimmer moves with velocity 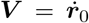, and angular velocity **Ω**. Using the Oseen approximation for hydrodynamic interactions between beads near a large sphere [31], we have:

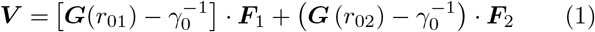

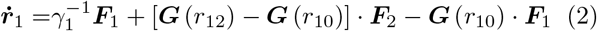

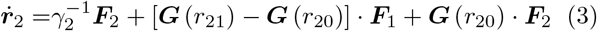

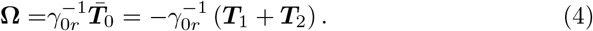

where ***r***_*ij*_ = ***r***_*i*_ − ***r***_*j*_, *γ*_*i*_ = 6*πηa*_*i*_, *γ*_0*r*_ = 8*πηa*_0_, 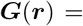 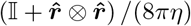, and dot denotes a time derivative. These reduce to a set of 10 equations for unknowns (***V***, **Ω**, 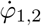, 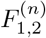) [31]. The method of quaternions was used to resolve singularities at *θ*_2_ = ±*π*/2 [32].

## Experiments

Our model aims to link the three-dimensional nature of flagellar beating to the cell’s free-swimming behaviour, but is *Chlamydomonas* flagellar beating truly non-planar, and if so, to what extent? A possible out-of-plane component was detected based on manual tracings of flagellar waveforms [16], but remains unclear. The same authors argued that helical swimming could result from transient asynchronies between the two flagella. Yet this is incompatible with more recent studies showing that asynchronies, or phase-slips, occur randomly, not periodically [26, 35]. We will use live-cell imaging and micromanipulation to prove that *Chlamydomonas* flagella beat with a persistent, non-planar pattern *in vivo*. To better visualise flagella motion, we aspirate cells onto pipettes to prevent body motion (following previous protocols [35, 36]). Cells are repositioned so that flagella are viewed (at 3000 fps) directly from above, with the eyespot delineating the *cis* (the flagellum closest to the eyespot) and *trans* flagella (Fig. 2a).

**FIG. 2.**
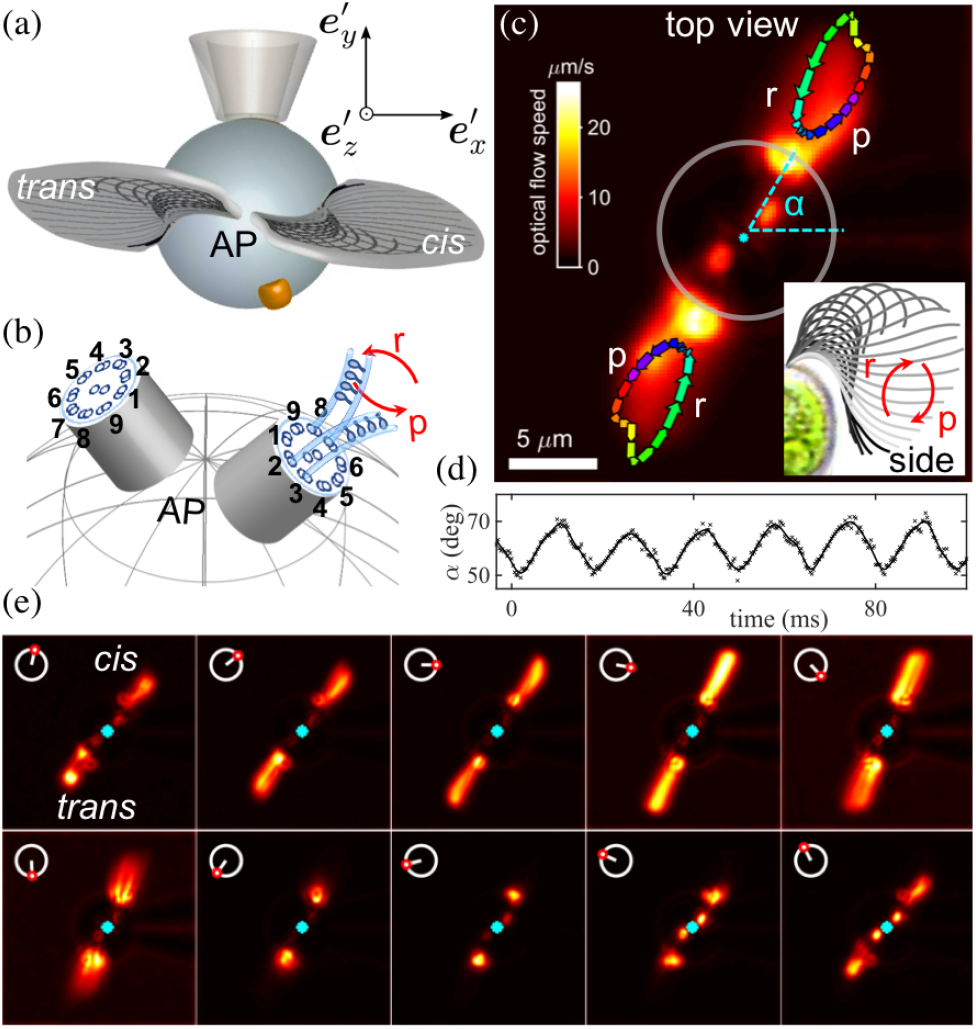
The non-planar beat pattern of *Chlamydomonas* flagella (see SM Video 1). (a) Cells are aspirated onto pipettes and imaged from the anterior pole (AP). An eyespot is located ~ 45° from the mean beat plane. (b) Basal bodies have a pre-defined symmetry with respect to axonemal microtubule doublets, numbered 1 → 9. Dyneins are separated into two groups, those on doublets operate 2, 3, 4 for the power (p) stroke, but 6, 7, 8 for the recovery (r) stroke [33]. (c) Waveforms are tracked by optic flow (arrows show direction of tip rotation), showing the relative offset between the basal bodies and the tilted beat planes. The beat plane (defined by *α*) rotates periodically (d). Waveforms are ordered by phase [34] and averaged over ~ 1000 consecutive cycles to show the phase-dependence of the non-planar beat pattern (e).

Our main findings are threefold. First, viewed from above the two basal bodies are displaced by a small clockwise offset, as reported previously in fixed specimens imaged using electron microscopy [37]. This offset is most evident when flagella motion is tracked by optical flow and overlaid to visualise the extent of movement (Fig. 2c). Second, the waveform is inherently non-planar. The *Chlamydomonas* power-recovery stroke cycle is usually imaged in the 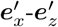 plane (Fig. 2c inset), here we reveal that there is also a significant component of motion in the 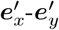 plane (Figure 2c-e, SM Video 1). During the power stroke, the flagella extend and pull away from the body while flagellar tips move in opposite directions; the direction of travel is immediately reversed during each subsequent recovery stroke. Over a stereotypical beat, the tips trace closed orbits (Fig. 2c). The rotary motion generates axial torques, balanced here by the micropipette. A freely-swimming cell, viewed from above, will therefore rotate clockwise (CW) during the power stroke but counter-clockwise (CCW) during the recovery stroke. This is analogous to the “two steps forward, one step backward” interpretation for the in-plane breaststroke, which arises due to flagella drag anisotropy. Third, the rotary motion is periodic (coincident with beat frequency), and is stable over thousands of cycles. To estimate the effective orbital tilt angle *β* from the high-speed videos, we use the following trignometric expression 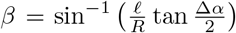, where Δ*α* is the angle subtended by the flagellum’s uppermost tip in the transverse 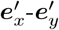 plane [31]. For *N* = 6 distinct cells, ~ 10^3^ consecutive beat cycles per individual, *β* = 0.3 rad ≈ 17.2° (see summary statistics in Table 1 [31]).

## Simulations

We input this estimate of *β* into our 3-bead model with cell-realistic parameters to determine if beat non-planarity can indeed generate axial rotation and helical swimming. All lengths are non-dimensionalised by *ℓ* (= 10 *μm*, typical cell size), forces by the average tangential flagellar force *F*_0_ (= 30 pN, typical force produced by a flagellum [38]), and *η* by 10^−3^ pN*μm*^−2^ (viscosity of water). When *β* = 0 (no tilt), pitch and roll (*θ*_2,3_) are suppressed, and our model reduces to the purely planar case investigated by other authors [24, 39]. This is when the only non-vanishing component of torque is along 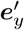 (equation (4)) and swimming is constrained to the cell’s frontal plane (Ω_*y*_, yaw).

For a non-planar beat (*β* ≠ 0), there is a non-vanishing resultant axial torque *T*_*z*_. When flagellar driving forces are equal 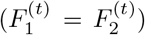, we have purely axial rotation, since other components of **Ω** cancel by symmetry. The resulting trajectory is linear, with **Ω** ∥ ***V*** . The trajectory can be a non-degenerate helix only if **Ω** ∦ ***V*** . Rotational symmetry about 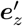 can be broken if (1) the scaffold shape is asymmetric; (2) flagella planes have different tilt (*β*_1_ ≠ *β*_2_); (3) flagella experience different driving forces 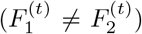. The first two cases sometimes produced trajectories that were irregular and non-helical [31]. Since we could not detect any obvious morphological asymmetries between the two *Chlamydomonas* flagella, we focus on case 2, and only consider asymmetric force profiles of the form 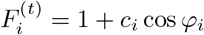 [30, 40].

For *β* > 0, symmetry-breaking introduced by hydrodynamic interactions between the beads results in either forward-CW or backward-CCW swimming at different phases of the beat cycle. The cycle-averaged swimming speed is given by 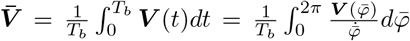 with flagellar period *T*_*b*_ and 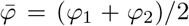, with an analogous expression for angular velocity 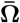 [31]. The dynamical system described by equations (2)–(3) is highly sensitive to swimmer shape. The strength of the hydrodynamic interactions between flagella beads are dictated by scaffold parameters, which determine net swimming (and rotation) direction per cycle, as in the 2D case [25, 40]. We choose configurations where 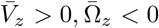 (CCW is positive), and average flagella beat frequencies *f*_*b*_ and axial rotation 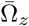 fall within experimental values.

We simulate free-swimming trajectories (Fig. 3) for a fixed scaffold shape (*a*_0_ = 0.53, *h* = 1.30, *R* = 0.60, *a* = 0.05), tilt angle *β* = 0.3, but different values of force asymmetry Δ*F* = *c*_2_ − *c*_1_. Fig. 3a is for purely axial rotation (Δ*F* = 0). When Δ*F* ≠ 0, trajectories are *superhelical*. Superhelices emerge as general solutions of all Low-*Re* swimming dynamics driven by periodic deformations [11], and have been observed in sperm swimming and in biaxial self-propelled particles under external torques [41]. A 1% difference in Δ*F* suffices to generate realistic superhelical trajectories (Fig. 3b). Here, the average trajectory of the centroid **r_0_** prescribes an outer helix with a pitch of ~ 95 *μm* and radius ~ 5.9*μm*, while “fast” helical swirls appear on the time scale of the flagellar period 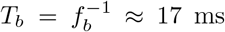. Less regular helices emerge with higher asymmetry (Fig. 3c,d). Frequencies of mean flagellar bead rotation *f*_*b*_ and axial rotation Ω_*z*_ vary with increasing orbital tilt angle *β* ∈ (0, 2*π*) (Fig. 3e). As expected, axial rotation velocity increases with beat non-planarity (higher *β*). A tilt angle *β* ≈ 0.3 corresponds to a rotation frequency of 1-2 Hz. We conclude that realistic helical swimming with a symmetric scaffold shape can be achieved using a very small force asymmetry between flagella, together with a small orbital tilt.

**FIG. 3.**
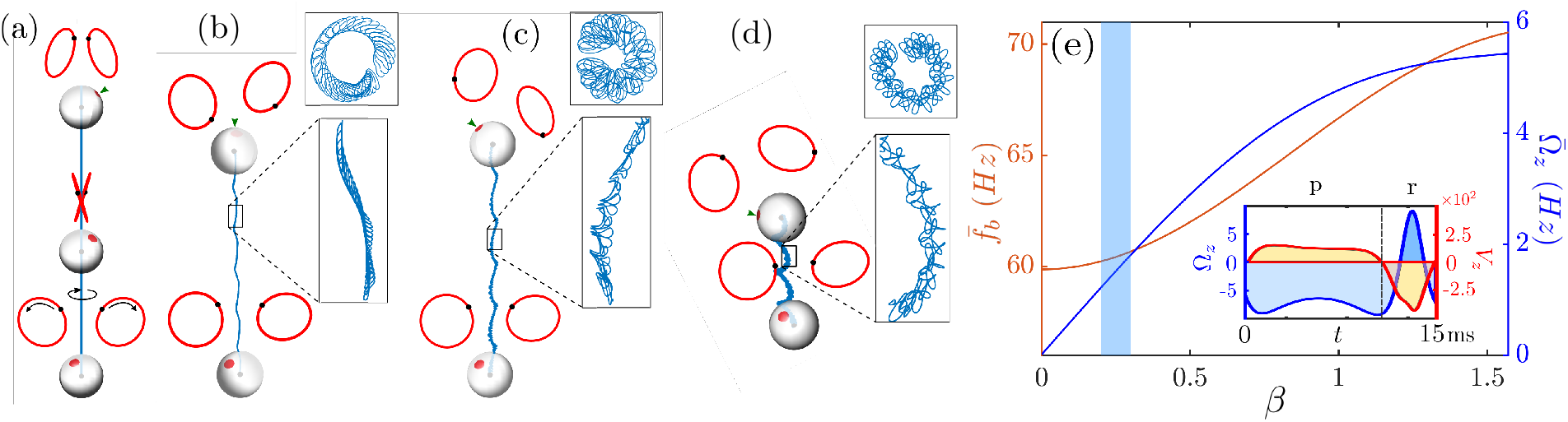
Non-planar flagellar beating leads to superhelical trajectories (eyespot: small arrowhead). For the same geometry *a*_0_ = 0.53, *h* = 1.30, *R* = 0.60, *β* = 0.30, *a* = 0.05, examples show (a) purely axial-rotation when *c*_1,2_ = 0.7, (b) ‘smooth’ superhelix for *c*_1_ = 0.7, *c*_2_ = 0.71, c) a superhelix with irregular inner helix for asymmetrical driving forces *c*_1_ = 0.6, *c*_2_ = 0.62, (d) a more irregular trajectory for *c*_1_ = 0.3, *c*_2_ = 0.5. (e) Mean beat frequency of the two flagella (orange) and axial component of angular velocity (blue) as functions of *β* (numerical data fitted with sum of sines). The shaded region marks range of *β* measured in experiments [31]. Inset: linear and angular velocities (in *μ*m/s and Hz) over one beat, note the CW-forward motion during the power (p) stoke and CCW-backward motion during the recovery (r) stroke.

## Phototactic steering

Can cells exploit asymmetric flagellar driving for trajectory control? We hypothesize that this mechanism underlies phototaxis - directed movement toward/away from light. *Chlamydomonas* cells perform positive or negative phototaxis, depending on the nature of the stimulus [19, 42]. Phototactic steering is associated with changes in both beat amplitude and frequency and is fine-tuned to the body rotation frequency [16, 20, 43]. In general, steering is accomplished by changing **Ω** [10, 44], which in turn changes helix properties (radius, pitch, orientation). Here, **Ω** and *f*_*b*_ are strongly coupled. To simulate phototaxis (Fig. 4), we assume that a light stimulus **I** either attenuates or accentuates Δ*F* depending on the alignment between the eyespot and stimulus [45]. Denoting by 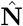 the vector normal to the eyespot surface, the intensity incident on the eyespot is *I*(*t*) = *I*_0_(*t*) cos *ϕ*(*t*) *H*(cos *ϕ*(*t*)), where 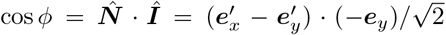, and *H* is the Heaviside step function. This framework can be applied to any type of taxis in response to a vectorial cue.

**FIG. 4.**
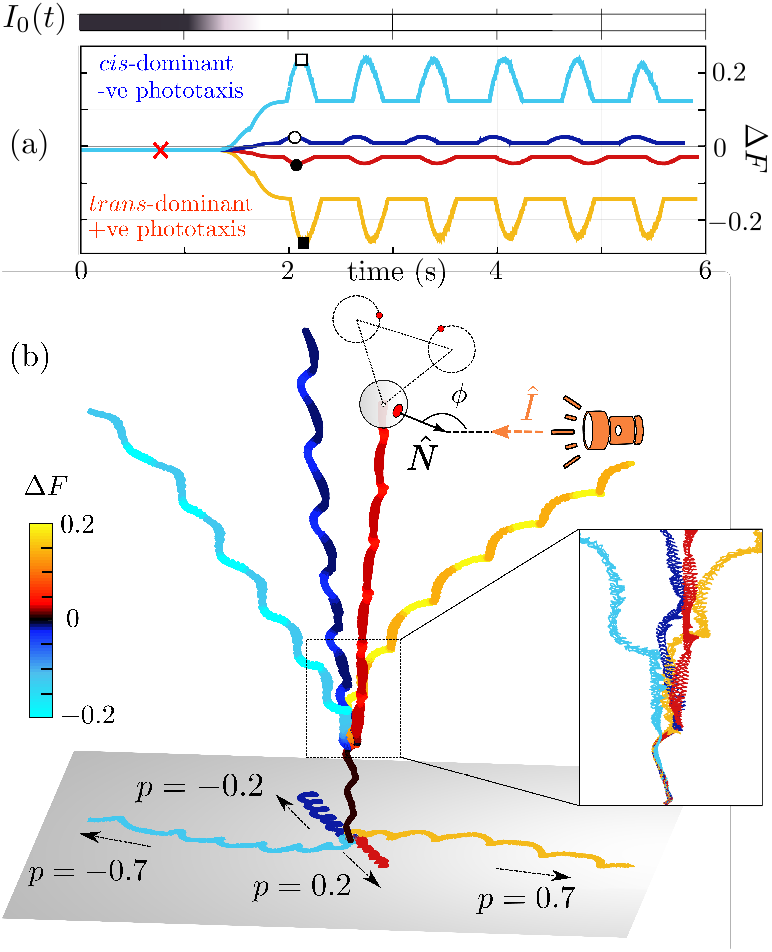
Flagellar dominance controls phototactic steering.(a) Time-series of stimulus intensity *I*_0_ (grayscale = intensity) and the response Δ*F* for four different *p* values. Markers indicate characteristic *transcis* frequencies: ×: 61 − 63*Hz*, □: 58-75Hz, ▅: 78-51Hz, ○: 51-73Hz, ●: 53-69Hz. (b) The corresponding 3D tracks show either positive or negative phototaxis. Color indicates force asymmetry Δ*F*. In all cases, *a*_0_ = 0.53, *h* = 1.3, *ℓ* = 1, *R* = 0.6, *β* = 0.3, *a* = 0.05, *c*_*cis*_(*t* = 0) = 0.71, *c*_*trans*_(0) = 0.7.

Flagella identity is important. When Δ*F* = 0.01 (no signal), the stronger (dominant) flagellum is the *cis* flagellum 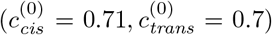. Force profiles are then modified to 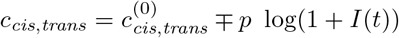, where *p* is the sensitivity of biflagellar dominance to the signal. Signs were chosen so that the model flagella responded differentially to the same signal [18], and in a direction compatible with experiments [46]. Importantly, here **Ω** emerges from the biflagellar force profiles, and was not assumed *a priori* to produce a regular helix. The resulting dynamics are characterised by a change in the pitch, radius and axis of the (super)helical trajectory. The sign of flagellar dominance, determines the sign of phototaxis. Compared to the no-light scenario, the cell turns towards the light when Δ*F* flips sign (*trans*-dominant), but away when the *cis*-flagellum becomes more dominant (Δ*F* more positive). This indicates that alignment to a vectorial stimulus is indeed possible simply by varying the two relative forcing profiles (Fig. 4).

## Concluding remarks

We showed for the first time that *Chlamydomonas* flagella beat with a persistent, 3D beat pattern *in vivo*, and quantified the extent of this non-planarity in terms of an orbital tilt. This complements a recent study which revealed that isolated axonemes, when reactivated with ATP, beat with a three-dimensional pattern [47]. It would seem that perfectly planar beats cannot be generated by an intrinsically chiral axoneme (Fig. 2b) [11, 48]. In *Chlamydomonas*, while the out-of-plane tilt may be fixed genetically, the in-plane beat pattern is dynamic [35]. Accounting for this 3D beat, we developed a three-bead hydrodynamic model with cell-realistic geometries which captures the sense and magnitude of axial rotation of real cells. Superhelical trajectories emerged directly from the flagellar motion, and were not prescribed. The model swims more slowly (15 – 30*μ*m/s) than live cells (50 – 100*μ*m/s [49]), suggesting limitations of a Stokeslet-type swimmer. Here, beat patterns/frequencies can only be changed by variable forcing, so that incorporating the full slender-filament dynamics will better mimic the amplitude-frequency coupling pertaining to real flagella. Basal coupling [50] may be additionally required to constrain biflagellar synchrony to enhance swimming and steering efficiency.

A small asymmetry (1%) in biflagellar driving forces was necessary and sufficient for helical (not just purely axial) swimming. Whether this is augmented by further structural asymmetries (if *β*_1_ ≠ *β*_2_) can be revealed by multifocal imaging, to measure the full torsional profile of each flagellum. Our simple three-bead swimmer was able to perform phototaxis, requiring only a stimulus-dependent source of asymmetry. Controllable steering is thus fairly insensitive to gait and the precise beating waveform. Indeed some algal biflagellates which do not use in-phase breaststrokes are nonetheless phototactic (e.g. *Polytoma*, *Chryptomonas*). Our work emphasizes the functional distinction between the two *Chlamydomonas* flagella, showing that a tunable and reversible biflagellar dominance likely operates in live cells for helical klinotaxis. Consistent with this view, mutant strains (*mia*, *bop*5-3, *ptx1*, and *lsp*1) have ‘smoother’ trajectories, and defective or weakened phototaxis [51–53]. Either flagellum can be dominant, depending on the stimulus. Indeed Ca^2+^ has been shown to produce opposite responses in reactivated *cis* and *trans* axonemes [18, 46]. Our results offer key insights into the physiological basis of dynamic sensorimotor control, as implemented in a simple, aneural organism with few morphological asymmetries. An important next step will be to determine how biflagellar dominance is regulated at the molecular level (e.g. of dynein activity).

We thank Ray Goldstein for financial support during the initial phase of this project, and Peter Ashwin and Gáspár Jékely for useful discussions. KYW is grateful for funding from an Academy of Medical Sciences Springboard Award, and the European Research Council (ERC) under the European Union’s Horizon 2020 research and innovation programme (grant no. 853560, *EvoMotion*).

