## Supplemental material (text) for "Control of helical navigation by three-dimensional flagellar beating"

Dario Cortese* and Kirsty Y. Wan†
*Living Systems Institute & College of Engineering, Mathematics and Physical Sciences*
*University of Exeter, Exeter EX4 4QD, United Kingdom*
(Dated: September 27, 2020)

We provide details of the single-cell experiments (section A) and the mathematical model (section B).

#### A: SINGLE-CELL MICROMANIPULATION EXPERIMENTS

Axenic cultures of *Chlamydomonas reinhardtii* (*Chlamydomonas* Center, CC125) were grown in exponential phase, to a density of $\sim 10^6$ cells per ml. Imaging and single-cell micromanipulation was conducted as previously described [1]. Additionally, upon pipette capture, cells were reoriented gently so that flagellar beating can be visualised directly from the anterior end (Figure 2a, main paper). Special care was taken to minimise cell mechanosensitive responses, and to ensure that no flagella were damaged during the reorientation procedure - any cells whose beat frequencies appeared unstable were duly discounted. See **SM Video 1** for a sample video (full length videos will be uploaded to a repository prior to publication). High-speed imaging (Phantom Vision Research) was conducted at 3000 frames per second. After each recording, an additional still image was taken with a Nikon DSLR camera (Nikon D300S) to determine the eyespot location (see Fig.1). This is important to distinguish the *cis* flagellum from the *trans* for each individual.

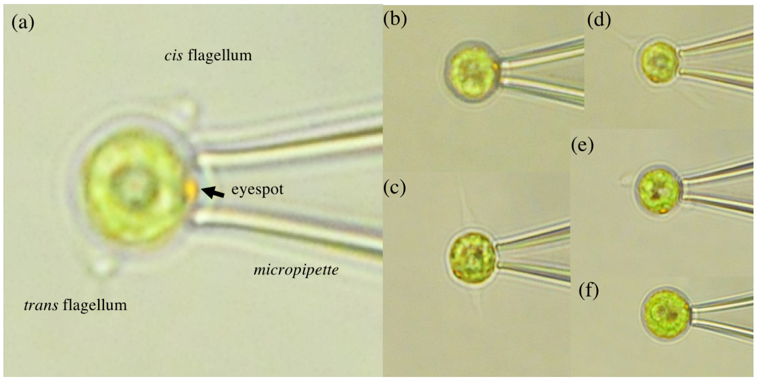


FIG. 1. Identifying the eyespot position from light-microscopy images. a)-f) represent 6 different individuals.

High-speed videos were analysed using custom image processing algorithms written in Matlab (Mathworks, R2018a). Briefly, a moving average subtraction was applied to the image stack, to highlight moving features (the flagella). The resulting grayscale images were then processed using the Horn-Schunck optical flow algorithm (Matlab Computer Vision Toolbox), and thresholded by optical flow magnitude to separate moving pixels (flagella) from the static background. The point cloud comprising all such 'moving' pixels from each frame was used to determine the spatial extent of flagellar movement. The speed of pixel 'flow' is a proxy for speed of movement of material points along the flagellum (Fig.2a).

The segmented region, corresponding to the boundary of the flagella pair, was used to determine an instantaneous flagella beat plane $\alpha$, on each frame. Oscillations in $\alpha$ were converted into a phase via the Hilbert transform (Fig.2b). The Fourier transform of $\alpha(t)$ was used to estimate the mean flagellar beat frequency over thousands of beats (Fig.2c), as well as the maximal angular deviation $\Delta\alpha$ between the start and end of each cycle (Fig.2d). Since $\Delta\alpha$ approximates the range of angles subtended by the flagella projected on the focal plane, we can use this to estimate the tilt angle $\beta$ according to $\beta = \sin^{-1}\left(\frac{\ell}{R}\tan\frac{\theta}{2}\right)$ (see Fig.3a for a graphical derivation). The equivalent geometric parameters $\ell$, $R$ vary from cell to cell. The anterior view allows us to estimate the scaffold geometry, namely the cell radius ($= 2a_0$), and the maximum distance from the centre of the cell body to the tip of the flagella ($\approx L + R$). We will further assume that $\ell \approx 2a_0$ - consistent with previously published *Chlamydomonas* waveforms (see also Fig.2, main text, inset). Cell geometry measurements were performed manually in Fiji [2].

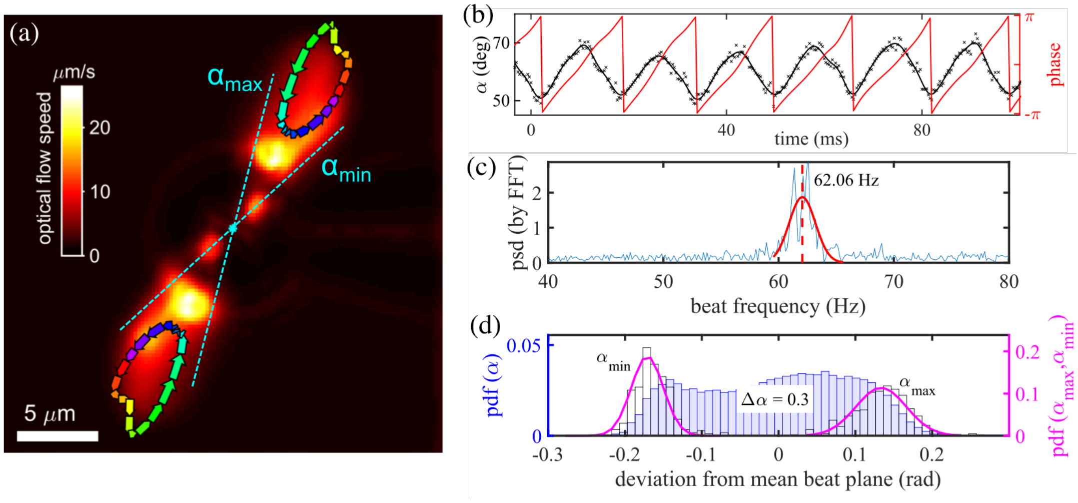


FIG. 2. Summary of video processing procedure. (a) Optical flow tracking was used to determine the position of the flagella. Flagellar tips trace nearly-closed orbits, following cyclic power and recovery strokes. (b) The instantaneous beat plane $\alpha(t)$ is used to define a phase, by Hilbert transform. The FFT of $\alpha(t)$ was used to estimate the mean beat frequency (c). (d) The angular deviation $\Delta\alpha$ is determined as the peak-to-peak separation between the distributions of the maximum ($\alpha_{\max}$) and minimum ($\alpha_{\min}$) orientation angles, respectively.

We summerize the results for 6 different cells, with 1000 beat cycles per individual. The results are summarised in Table I. From which we estimate $\beta \sim 0.3$.

Video frames were further ordered by Hilbert phase, and waveforms corresponding to the same phase (isophase waveforms) were grouped together and averaged (Fig.4). The average progression of the beat-plane with respect to Hilbert phase, is shown in (Fig.3b).

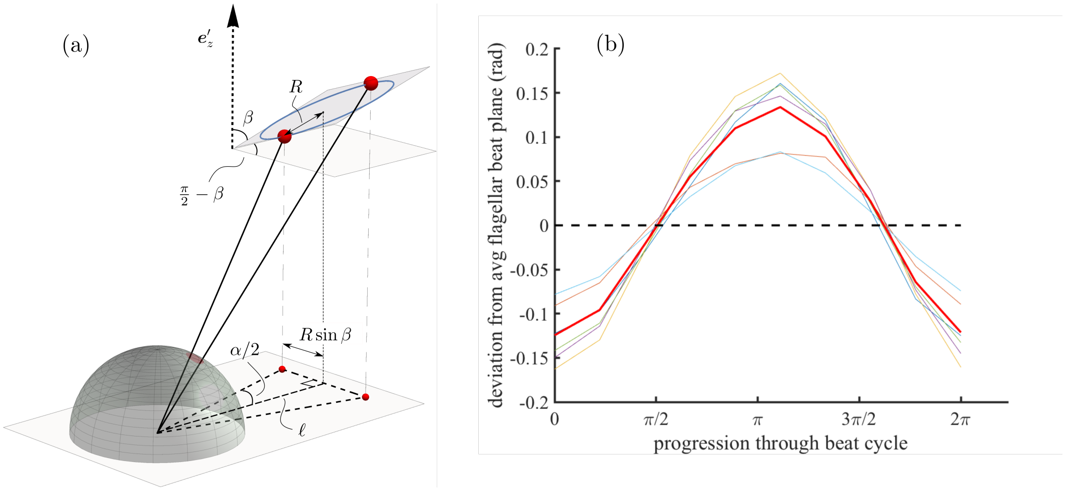


FIG. 3. (a) Relationship between the angle measured in the top-view experiments ($\alpha$), and the tilt angle between the flagellar plane and the posterior-anterior axis $\boldsymbol{e}'_z$. (b) Cyclic rotation of the beat plane as a function of Hilbert phase. Thick red line - average over all 6 individuals.

| Cell | No. Beats | Beat Frequency | Cell Diameter ($\mu m$) $\sim 2a_0$ | Tip Distance ($\mu m$) $\sim \ell + R$ | $\ell/R$ | $\Delta\alpha$ |
|---|---|---|---|---|---|---|
| 1 | 1074 | 61.7 | 6.8 | 12.5 | 1.2 | 0.22 |
| 2 | 708 | 61.5 | 11.7 | 16.5 | 2.4 | 0.25 |
| 3 | 293 | 63.1 | 9.7 | 13.4 | 2.6 | 0.23 |
| 4 | 609 | 61.5 | 7.9 | 13.6 | 1.4 | 0.27 |
| 5 | 333 | 58.5 | 8.8 | 12.8 | 2.2 | 0.27 |
| 6 | 424 | 62.1 | 9.2 | 13.2 | 2.3 | 0.30 |
| Average | | | 9.0 | 61.4 | 2.0 | 0.26 |

TABLE I. Parameter estimates from data. The measured scaffold geometries are consistent with simulations.

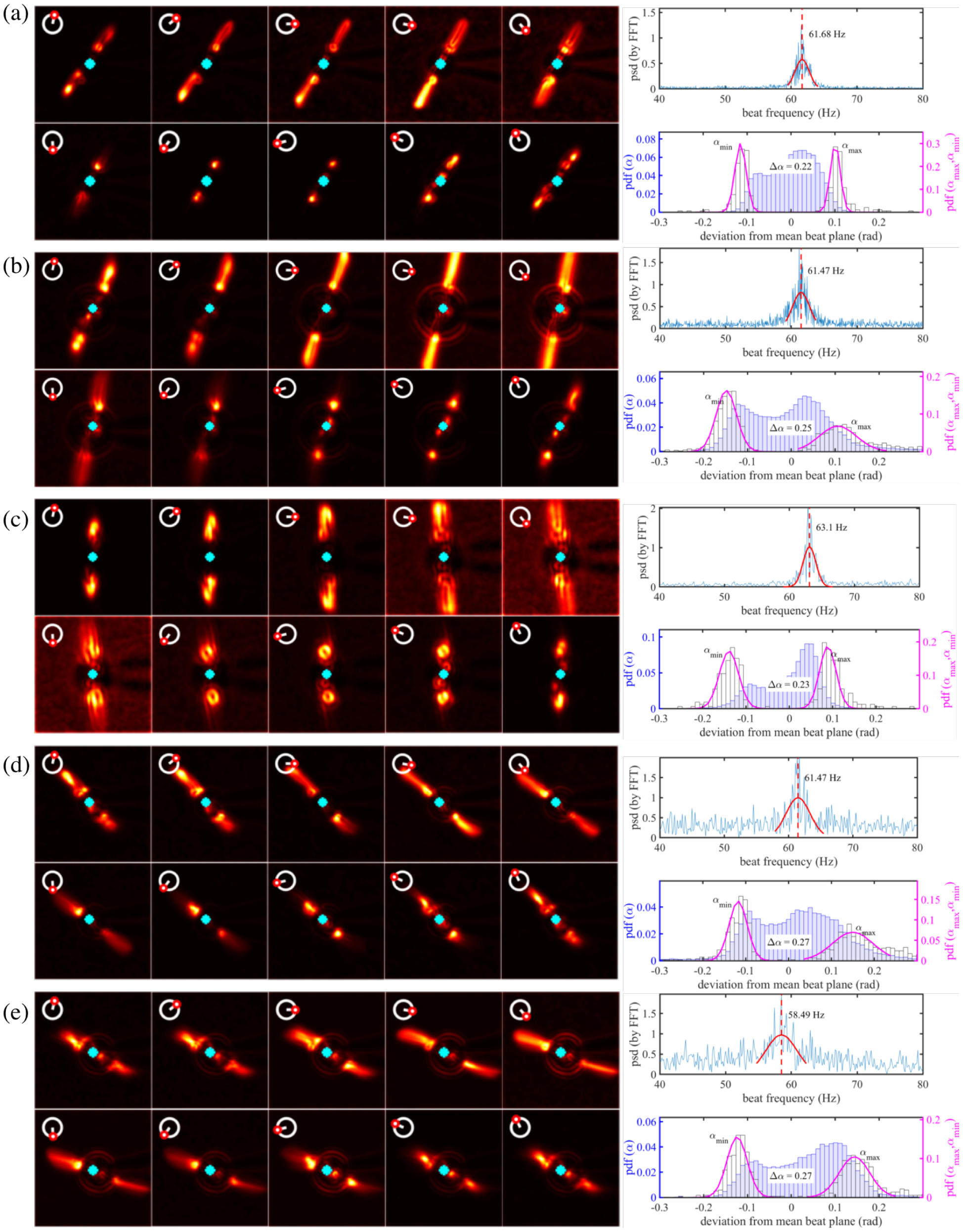


FIG. 4. Extracting 3D tilt angles from data. All cells show a consistent CCW-rotation of the beat plane during the power stroke, and CW-rotation during the subsequent recovery stroke. Axial torques are balanced by the pipette. A free-swimming cell will therefore rotate CW during the power stroke and CCW during the recovery stroke, analogous to the two-steps forward and one-step back motion in the swimming plane.

### B: DETAILS OF THE HYDRODYNAMIC MODEL

Please refer to the main text and Fig. 1 for model description and set-up.

#### Body frame of reference

We define a rotating frame of reference $\left(\boldsymbol{e}'_x, \boldsymbol{e}'_y, \boldsymbol{e}'_z\right)$ at rest with the body sphere, characterised by three Euler angles $\boldsymbol{\theta} = (\theta_1, \theta_2, \theta_3)$ and the following transformation rules: $\mathbf{e}'_i = \boldsymbol{L}(\boldsymbol{\theta})\,\mathbf{e}_i$. We adopt the Tait-Bryan angle convention (with a $y - x'' - z'$ or 3-1-2 intrinsic definition), where $\theta_1, \theta_2, \theta_3$ are denominated yaw, pitch and roll angles and describe the clockwise rotations around the $\boldsymbol{e}'_x, \boldsymbol{e}'_y, \boldsymbol{e}'_z$ axes, respectively. The matrix $\boldsymbol{L}(\boldsymbol{\theta}) = L_{ij}\boldsymbol{e}_i\boldsymbol{e}_j = \boldsymbol{R}_x(\theta_1)\,\boldsymbol{R}_y(\theta_2)\,\boldsymbol{R}_z(\theta_3)$, where

$$\boldsymbol{R}_x(\theta_1) = \begin{pmatrix} 1 & 0 & 0 \\ 0 & c_1 & -s_1 \\ 0 & s_1 & c_1 \end{pmatrix}, \quad \boldsymbol{R}_y(\theta_2) = \begin{pmatrix} c_2 & 0 & s_2 \\ 0 & 1 & 0 \\ -s_2 & 0 & c_2 \end{pmatrix}, \quad \boldsymbol{R}_z(\theta_3) = \begin{pmatrix} c_3 & -s_3 & 0 \\ s_3 & c_3 & 0 \\ 0 & 0 & 1 \end{pmatrix},$$

and $c_i = \cos\theta_i$ and $s_i = \sin\theta_i$. In this description [3],

$$\dot{\boldsymbol{\theta}} = \boldsymbol{E}^{-1}\boldsymbol{\omega}, \quad \boldsymbol{E} = \begin{pmatrix} 1 & 0 & s_2 \\ 0 & c_1 & -s_1c_2 \\ 0 & s_1 & c_1c_2 \end{pmatrix}$$

However, $\boldsymbol{E}^{-1}$ has a coordinate-type singularity usually referred to as *gimbal lock* at $\theta_2 = \pi/2$:

$$\boldsymbol{E}^{-1} = \frac{1}{c_2}\begin{pmatrix} 1 & s_1s_2 & -c_1s_2 \\ 0 & c_1c_2 & s_1c_2 \\ 0 & -s_1 & c_1 \end{pmatrix}.$$

To avoid this, we can adopt a quaternion fomulation [4]. We encode all the information regarding the orientation of the swimmer in the following unit quaternion

$$\mathbf{q}_r(\alpha, \boldsymbol{n}) := e^{\frac{\theta}{2}(n_x\boldsymbol{i}+n_y\boldsymbol{j}+n_z\boldsymbol{k})} = \begin{bmatrix} \cos\frac{\alpha}{2} \\ \boldsymbol{n}\sin\frac{\alpha}{2} \end{bmatrix}$$

Vectors are rotated according to the quaternion multiplication [4] and the orientation dynamics is obtained via the relationship between quaternion rates and angular velocity

$$\dot{\mathbf{q}} = \frac{1}{2}\mathbf{q} \star \begin{bmatrix} 0 \\ \boldsymbol{\omega} \end{bmatrix} = \frac{1}{2}\begin{pmatrix} 0 & -\omega_1 & -\omega_2 & -\omega_3 \\ \omega_1 & 0 & \omega_3 & -\omega_2 \\ \omega_2 & -\omega_3 & 0 & \omega_1 \\ \omega_3 & \omega_2 & -\omega_1 & 0 \end{pmatrix}\mathbf{q} \tag{1}$$

Under the assumption of constant $\boldsymbol{\omega}$ direction, which is satisfied for an interval $\Delta t$ such that $\boldsymbol{\omega}(t+\Delta t)/\left|\boldsymbol{\omega}(t+\Delta t)\right| \simeq \boldsymbol{\omega}(t)/\left|\boldsymbol{\omega}(t)\right|$, the solution to (1) is

$$\mathbf{q}(t) \simeq \exp\left(\frac{1}{2}\boldsymbol{\omega}\Delta t\right) \star \mathbf{q}_0.$$

From this expression, the Euler angles can be recovered using the following formulae

$$\boldsymbol{\theta} = \begin{pmatrix} \arctan\left(2\frac{q_0q_3+q_1q_2}{1-2\left(q_2^2+q_3^2\right)}\right) \\ \arcsin\left(2q_0q_2 - 2q_1q_3\right) \\ \arctan\left(2\frac{q_0q_1+q_2q_3}{1-2\left(q_1^2+q_2^2\right)}\right) \end{pmatrix}$$

#### Shape of the swimmer

The center of the two flagellar beads' orbits are $\boldsymbol{s}_i = \mathbf{r}_0 + (-1)^i \, \ell \boldsymbol{e}'_x + h\boldsymbol{e}'_z, \; i = 1, 2$, while the position of the two flagellar beads is given by the vector expression

$$\mathbf{r}_i = \boldsymbol{s}_i \left(\mathbf{r}_0, \boldsymbol{\theta}\right) + R\hat{\boldsymbol{n}}_i \left(\boldsymbol{\theta}, \varphi_i, \beta\right), \quad i = 1, 2$$

where $\hat{\boldsymbol{n}}_i$ is the normal vector that connects $\boldsymbol{s}_i$ to the moving bead, and thus contains information about the phase $\varphi_i$ of the bead along its orbit as well as the orientation $\boldsymbol{\theta}$ of the latter.

We study a 3D configuration in which the orbit of the flagellar beads lies on a plane $\pi_{(\beta)}$, obtained by applying a rotation of angle $\beta$ around $\boldsymbol{e}'_x$ . In this case we have

$$\hat{\boldsymbol{n}}_i = (-1)^{i+1} \cos\varphi_i \boldsymbol{e}'_x + (-1)^{i+1} \sin\beta \sin\varphi_i \boldsymbol{e}'_y + \cos\beta \sin\varphi_i \boldsymbol{e}'_z.$$

The plane $\pi_{(\beta)}$ is fixed with respect to the swimmer. The system has therefore 8 degrees of freedom: $(x_0, y_0, z_0, \theta_1, \theta_2, \theta_3, \varphi_1, \varphi_2)$, and we keep $h, \ell, \alpha, \beta, R, a, b$ as parameters. The translational velocities of the three beads are found by differentiation:

$$\dot{\boldsymbol{r}}_i = \dot{\boldsymbol{r}}_0 + \boldsymbol{\omega} \times \mathbf{r}_i + \dot{\mathbf{r}}'_i = \dot{\boldsymbol{r}}_0 + \boldsymbol{\omega} \times \boldsymbol{r}_i + R\dot{\varphi}_i$$

#### Hydrodynamic forces

The driving force on each flagellum is the tangential component $F_i^{(t)}\hat{\mathbf{t}}_i$ and it is balanced by tangential component of the local hydrodynamic drag (given by Stokes law), $\mathbf{F}_i + \mathbf{F}_i^{(\mathrm{drag})} = \mathbf{0}$. $\mathbf{F}_i^{(\mathrm{drag})}$ is linear in the relative flow velocity:

$$\mathbf{F}_i^{(\mathrm{drag})} \cdot \hat{\mathbf{t}}_i = -\gamma_i \hat{\mathbf{t}}_i \cdot \left[\dot{\mathbf{r}}_i - \mathbf{u}\left(\mathbf{r}_i\right)\right], \quad \gamma_i = 6\pi\eta a_i$$

where $\mathbf{u}$ is the fluid flow. Because each bead is constrained to move on a circular trajectory (in the frame of the swimmer), the tangential force balance becomes

$$\dot{\mathbf{r}}_i \cdot \hat{\mathbf{t}}_i = \gamma_i^{-1} F_i^{(t)} + \hat{\mathbf{t}}_i \cdot \mathbf{u}\left(\mathbf{r}_i\right) \tag{2}$$

Similar relationships apply to the rotational motion of each bead:

$$\bar{\mathbf{T}}_i - \gamma_{r,i} \boldsymbol{\Omega}_i = \mathbf{0}, \quad \gamma_i^{(r)} = 8\pi\eta a$$

The local fluid flow $\mathbf{u}(\mathbf{r}_i)$ is perturbed by the motion of the other flagella, and this effect mediates the interaction among different beads. In the zero Reynolds number approximation, these interactions (*i.e.* the fluid flow produced at $\mathbf{r}_i$ because of a force being applied at $\mathbf{r}_j$) can be expressed in terms of *linear* relationships between forces and velocities:

$$\mathbf{u}\left(\mathbf{r}_i\right) = \sum_{j \neq i} \mathbf{H}\left(\mathbf{r}_i, \mathbf{r}_j\right) \cdot \mathbf{F}_j$$

where $\mathbf{H}$ is a two-rank tensor that models the interaction. This can be incorporated in (2), to obtain

$$\dot{\mathbf{r}}_i = \gamma_i^{-1} \boldsymbol{F}_i + \sum_{j \neq i} \mathbf{H}\left(\mathbf{r}_i, \mathbf{r}_j\right) \cdot \boldsymbol{F}_j.$$

Using the formalism of Reichert [5], we can write the general relationship between velocities, forces and torques as

$$\dot{\mathbf{r}}_i = \sum_j \left(\mathbf{M}_{ij}^{tt} \cdot \mathbf{F}_j + \mathbf{M}_{ij}^{tr} \cdot \bar{\mathbf{T}}_j\right), \quad \boldsymbol{\Omega}_i = \sum_j \left(\mathbf{M}_{ij}^{rt} \cdot \mathbf{F}_j + \mathbf{M}_{ij}^{rr} \cdot \bar{\mathbf{T}}_j\right) \tag{3}$$

where $\mathbf{M}$ is the mobility matrix, the superscripts $r, t$ stand for rotation and translation, respectively. The explicit form of $\mathbf{M}$ depends on $\gamma_i, \gamma_{r,i}$ $\boldsymbol{H}$ and on how we model the hydrodynamic interactions among the beads.

*Three Stokeslets: point force approximation*

If we consider each sphere in the far field of the other, the geometrical details of the configuration are significantly simplified. In this case, we can model the interaction between two individual beads by using a multipole expansion and neglecting high order terms in the separation $\mathbf{r}_{ij} = \mathbf{r}_i - \mathbf{r}_j$, with $i, j = 0, 1, 2$. The first order term of this expansion corresponds to approximating the stresses exerted by the bead on the fluid with a point force, and $\boldsymbol{H}$ can be expressed as follows

$$\boldsymbol{H}(\mathbf{r}_i, \mathbf{r}_j) = \mathbf{G}(\mathbf{r}_{ij}), \quad \mathbf{G}(\mathbf{r}) = \frac{1}{8\pi\eta r}\left(\mathbb{I} + \hat{\boldsymbol{r}} \otimes \hat{\boldsymbol{r}}\right)$$

$\mathbf{G}(\mathbf{r})$ is commonly referred to as the Oseen's tensor or *stokeslet.* Note that the spheres are point-like particles, therefore their respective radii, as well as any rotational effects, do not appear in this approximation of the hydrodynamic interaction. However, the radius appears in the viscous friction coefficient $\gamma$ that models the interaction between each sphere and the local fluid flow. This approach does include far field interactions, but completely neglects finite size and therefore the rotational part of the hydrodynamic stresses. We can express the relationship between velocities and forces as

$$\begin{aligned}
\dot{\boldsymbol{r}}_0 &= \left[\mathbf{G}(r_{01}) - \gamma_0^{-1}\right] \cdot \boldsymbol{F}_1 + \left(\mathbf{G}(r_{02}) - \gamma_0^{-1}\right) \cdot \boldsymbol{F}_2 \\
\dot{\boldsymbol{r}}_1 &= \gamma_1^{-1}\boldsymbol{F}_1 + \left[\mathbf{G}(r_{12}) - \mathbf{G}(r_{10})\right] \cdot \boldsymbol{F}_2 - \mathbf{G}(r_{10}) \cdot \boldsymbol{F}_1 \\
\dot{\mathbf{r}}_2 &= \gamma_2^{-1}\boldsymbol{F}_2 + \left[\mathbf{G}(r_{21}) - \mathbf{G}(r_{20})\right] \cdot \boldsymbol{F}_1 + \mathbf{G}(r_{20}) \cdot \boldsymbol{F}_2
\end{aligned} \tag{4}$$

where we have used the force-free condition (7), and $\gamma_0 = 6\pi\eta b, \quad \gamma_{1,2} = 6\pi\eta a.$

*Two Stokeslets near a spherical body*

A more accurate description of the interaction among the flagella and the body of the microorganism can be achieved as follows. The body can be modelled as a large sphere of radius $A \gg a$, whereas the two flagella are still described in terms of small sphere that interact with the fluid as point forces. Considering the body as an extended spherical solid object introduces more detail in the interaction of each flagellum with the cellular wall. The corresponding modified Green function was derived by Oseen and details of its derivation are found in [6–8].

**Force and torque-free condition**

The system is not subject to any external forces or torques, therefore we can impose the force-free and torque-free conditions

$$\boldsymbol{F} = \boldsymbol{F}_0 + \boldsymbol{F}_1 + \boldsymbol{F}_2 = \mathbf{0} \tag{5}$$

$$\boldsymbol{T} = \boldsymbol{T}_0 + \boldsymbol{T}_1 + \boldsymbol{T}_2 = -\mathbf{r}_0 \times (\boldsymbol{F}_1 + \boldsymbol{F}_2) + \mathbf{r}_1 \times \boldsymbol{F}_1 + \mathbf{r}_2 \times \boldsymbol{F}_2 + \bar{\boldsymbol{T}}_0 + \bar{\boldsymbol{T}}_1 + \bar{\boldsymbol{T}}_2 = \mathbf{0}$$

where $\bar{\boldsymbol{T}}_i$ is the intrinsic torque due to the rotation of the $i$-th bead around an axes passing through its center.

In the following, we will discuss two geometrical limits. The first consists of assuming all the beads have radii $a, b \ll \ell$, so that they act as point forces, and their intrinsic rotation is negligible, so that we can neglect the intrinsic torques $\bar{\boldsymbol{T}}_i$. In the second limit, adopted by other authors [9], we assume that only the flagellar beads satisfy $a \ll b \ll \ell$, and therefore the intrinsic torque of the system can be reduced to $\bar{\boldsymbol{T}}_0 \gg \bar{\boldsymbol{T}}_{1,2}$. Note that this description is similar to a previous model [10] in that it allows the body to spin around its axis, but it is different because it considers only the body bead's rotations; furthermore, our model is full three-dimensional, and this makes the torque acting on $\boldsymbol{T}_0$ are more complicated object, as it can have any direction in $\mathbb{R}^3$. In the simulations referred to in the main manuscript, the second limit was implemented.

Therefore, to allow for the two limits to be studied we write the torque free condition as

$$\bar{\boldsymbol{T}}_0 + (\mathbf{r}_1 - \mathbf{r}_0) \times \boldsymbol{F}_1 + (\mathbf{r}_2 - \mathbf{r}_0) \times \boldsymbol{F}_2 = \mathbf{0} \tag{6}$$

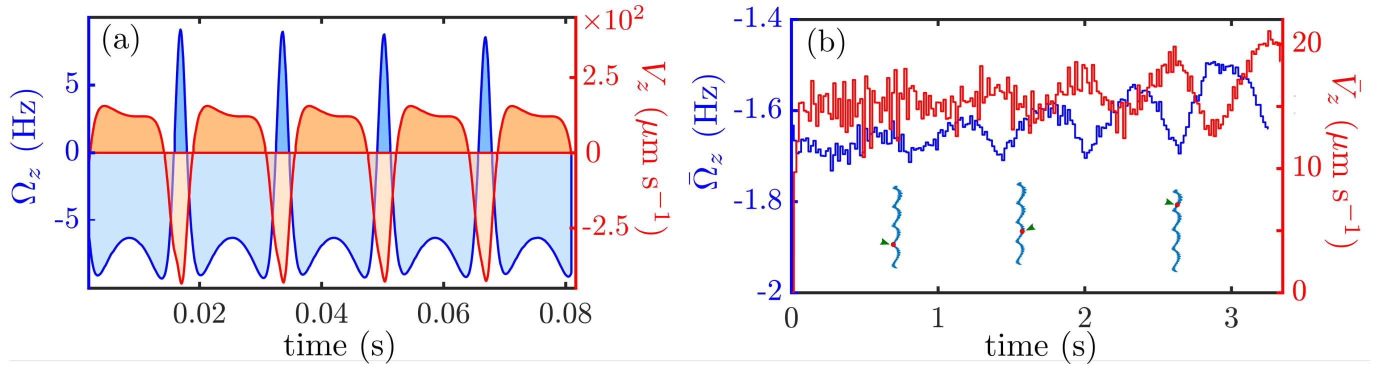


FIG. 5. Top: time evolution of the axial component of $\Omega$ and the forward velocity $V_z$.The graph shows 5 full flagellar beats, characterised by alternating forward-backward and clockwise-anticlockwise phases, corresponding to the power and recovery strokes. The integral quantities $\bar{V}_z$ and $\bar{\Omega}_z$, are directly proportional to the area under each of the curves. Bottom: $\bar{V}_z$ and $\bar{\Omega}_z$ vary as the cell swims on its helical trajectory. The diagrams under the curves indicate the position of the cell body relative to the superhelical track. Geometry $a_0 = 0.53,\ h = 1.30,\ R = 0.60,\ \beta = 0.30,\ a = 0.05$, smooth superhelical trajectory with $c_1 = 0.7,\ c_2 = 0.71$

The flagellar beads are driven by the tangential component of the respective driving force, which is in principle phase-dependent. The force-free condition reads:

$$\boldsymbol{F}_i = F_i^{(t)}(\varphi_i)\,\hat{\boldsymbol{t}}_i + F_i^{(n)}\hat{\boldsymbol{n}}_i \tag{7}$$

Once the tangential driving forces $F_{1,2}^{(t)}$ are known, (6-7) consist of 6 equations in 5 unknowns: $\boldsymbol{F}_0, F_{1,2}^{(n)}$.

*The role of the intrinsic hydrodynamic torque*

If we consider three point forces when modelling the interactions between the solid beads, the torque balance (6) reads simply $(\mathbf{r}_1 - \mathbf{r}_0) \times \boldsymbol{F}_1 + (\mathbf{r}_2 - \mathbf{r}_0) \times \boldsymbol{F}_2 = \mathbf{0}$, and the only torques that appear in this equation are due to the motion of the beads in the frame of reference of the laboratory. However, if we model the flagella as point forces and the body of the swimmer as a solid sphere of radius $A \gg a$, we have to include the intrinsic hydrodynamic torque $\bar{\mathbf{T}}_0$ due to the rotation of the body. This torque, thanks to the linearity of the Stokes equation, is linearly linked to the components of the angular velocity of the body $\boldsymbol{\omega}_0$:

$$\boldsymbol{\omega}_0 = \mathbf{M}_{ij}^{rt} \cdot \mathbf{F}_j + \mathbf{M}_{00}^{rr} \cdot \bar{\mathbf{T}}_0,$$

where we have used the fact that $\bar{\mathbf{T}}_{1,2} = \mathbf{0}$. For a solid sphere of radius $A$, it can be shown [5] that $\mathbf{M}_{ij}^{rt} = \mathbf{0}, \quad M_{00} = \gamma_{0r}^{-1} = (8\pi\eta A)^{-1}$. Since we are assuming here that the centre of the flagellar orbits are rigidly fixed with respect to the body sphere, we can conclude that $\boldsymbol{\omega}_0 = \boldsymbol{\omega}$, *i.e.* the intrinsic torque of the body sphere is directly related to the rotational velocity of the swimmer. This finally implies that $\boldsymbol{\omega} = (8\pi\eta A)^{-1}\,\bar{\mathbf{T}}_0$.

**Nondimensional equations and solution method**

We non-dimensionalise the governing equations by measuring quantities in units of the fluid viscosity $\eta \approx 10^{-3}$pN$\mu$m$^{-2}$s, the swimmer's length scale $\ell \approx 10\mu$m, and the typical average flagellar force $F_0 \approx 30$pN [11].

In total, we have an algebraic system of 10 equations for 10 unknowns $\left(\dot{\mathbf{r}}_0, \boldsymbol{\omega}, \dot{\varphi}_{1,2}, F_{1,2}^{(n)}\right)$.

$\boldsymbol{V} = \dot{\mathbf{r}}_0$ and $\boldsymbol{\omega}$ are the instantaneous rotational and linear velocities of the cell body. Because of the periodic nature of the flagellar motion encoded in $\varphi_{1,2}$, these oscillate between positive (forward, clock-wise) and negative (backward, anti-clockwise) values. This alternation is represented in Fig.5 and the inset of Fig.3(e) in the main manuscript, where the asymmetry between the power and recovery stroke is evident within each beat cycle. The net velocities are proportional to the area (with sign) under the graph in Fig.5, obtained by integrating the instantaneous expressions according to the following formulae:

$$\bar{\boldsymbol{V}} = \frac{1}{T_b}\int_0^{T_b} \boldsymbol{V}(t)dt = \frac{1}{T_b}\int_0^{2\pi} \frac{\boldsymbol{V}}{\dot{\phi}}d\phi, \quad \bar{\boldsymbol{\Omega}} = \frac{1}{T_b}\int_0^{T_b} \boldsymbol{\Omega}(t)dt = \frac{1}{T_b}\int_0^{2\pi} \frac{\boldsymbol{\Omega}}{\dot{\phi}}d\phi$$

$$\phi = \varphi_1 - \varphi_2, \quad T_b = \int_0^{T_b} dt = \int_0^{2\pi} \frac{d\phi}{\dot{\phi}}$$

**Numerical methods**

We adopted an adaptive time-stepping strategy. At each simulation step $i$, the velocities and resulting positions $\boldsymbol{X}$ are calculated for a time step $\Delta t$ and its half $\Delta t/2$. An error is calculated by evaluating the difference between the phases values in the two cases

$$\epsilon = \max_i \left[\varphi_i^{(\Delta t/2)} - \varphi_i^{(\Delta t)}\right].$$

If $\epsilon$ is larger than a predetermined threshold $\epsilon_{max}$, the time step is halved: $\Delta t \to \Delta t/2$. On the other hand, if $\epsilon < \epsilon_{max}$, the velocities and positions are calculated as

$$\boldsymbol{X_i} = 2\boldsymbol{X}_i^{(\Delta t/2)} - \boldsymbol{X}_i^{(\Delta t)}.$$

After $N_a$ accurate steps (for which $\epsilon < \epsilon_{max}$), the time st ep $\Delta t$ is incremented by a factor $g$:

$$\Delta t \to \Delta t(1 + gN_a).$$

In this work, we have used $N_a = 5 - 10$, $\epsilon_{max} = 0.05 - 0.2$, $g = 0.02$.

---

*
†

[1] K. Y. Wan, K. C. Leptos, and R. E. Goldstein, Lag, lock, sync, slip: the many 'phases' of coupled flagella, Journal of the Royal Society Interface **11**, 20131160 (2014).
[2] J. Schindelin, I. Arganda-Carreras, E. Frise, V. Kaynig, M. Longair, T. Pietzsch, S. Preibisch, C. Rueden, S. Saalfeld, B. Schmid, *et al.*, Fiji: an open-source platform for biological-image analysis, Nature methods **9**, 676 (2012).
[3] M. Boyle, The Integration of Angular Velocity, Adv. Appl. Clifford Algebras **27**, 2345 (2017).
[4] J. Diebel, Representing attitude: Euler angles, unit quaternions, and rotation vectors, Matrix **58**, 1 (2006).
[5] M. Reichert, Hydrodynamic interactions in colloidal and biological systems, , 197.
[6] C. Maul and S. Kim, Image of a point force in a spherical container and its connection to the Lorentz reflection formula, J Eng Math **30**, 119 (1996).
[7] S. E. Spagnolie, G. R. Moreno-Flores, D. Bartolo, and E. Lauga, Geometric capture and escape of a microswimmer colliding with an obstacle, Soft Matter **11**, 3396 (2015).
[8] J. J. L. Higdon, A hydrodynamic analysis of flagellar propulsion, Journal of Fluid Mechanics **90**, 685 (1979).
[9] R. R. Bennett and R. Golestanian, Emergent Run-and-Tumble Behavior in a Simple Model of Chlamydomonas with Intrinsic Noise, Phys. Rev. Lett. **110**, 148102 (2013).
[10] K. Polotzek and B. M. Friedrich, A three-sphere swimmer for flagellar synchronization, New Journal of Physics **15**, 045005 (2013).
[11] T. J. Böddeker, S. Karpitschka, C. T. Kreis, Q. Magdelaine, and O. Bäumchen, Dynamic force measurements on swimming chlamydomonas cells using micropipette force sensors, Journal of the Royal Society Interface **17**, 20190580 (2020).
